# Use of a novel knock-in allele of *Pkd1* identifies nicotinamide nucleotide dehydrogenase as a mitochondrial binding partner of polycystin-1

**DOI:** 10.1101/2022.04.08.487705

**Authors:** Cheng-Chao Lin, Luis F. Menezes, Elisabeth Pearson, Fang Zhou, Yu Ishimoto, D. Eric Anderson, Gregory G. Germino

## Abstract

The localization and function of Polycystin-1, the protein encoded by the gene most commonly mutated in autosomal dominant polycystic kidney disease, remains controversial. We have recently reported that its C-terminus is cleaved and traffics to the mitochondria rather than to the nucleus as had been previously described, and we found that absence of PC1 resulted in fragmented mitochondrial networks and increased mitochondrial membrane potential. Direct visualization of PC1 in mitochondria was only possible, however, after over-expression of recombinant, fluorescently labeled-PC1 in a cell culture system. To resolve the issue, we generated a new mouse model with three copies of the HA epitope and eGFP knocked-in frame into the endogenous mouse *Pkd1* gene by CRISPR/Cas9. We show that the modified allele is fully functional but the eGFP-tagged protein cannot be detected without antibody amplification methods. We were, however, able to use nanobody-coupled beads and large quantities of tissue to isolate a PC1 interactome and verify nicotinamide nucleotide transhydrogenase (Nnt) as a mitochondrial partner, linking PC1 to regulation of reactive oxygen species levels in the mitochondria. Loss of *Nnt* function had no significant effect on renal cystic disease in *Pkd1* mutants but treatment of young mice with early onset cystic disease with n-acetyl-cysteine (NAC) provided modest benefit only in the Nnt^+/+^ genetic background. These studies suggest that new methods and brighter tags will be required to track endogenous PC1, but this new mouse model will be a valuable resource for characterizing the protein interactome of endogenous PC1. The data also support our prior findings that the PC1 C-terminus localizes to mitochondria and regulates their function.

## Introduction

The gene most commonly mutated in autosomal dominant polycystic kidney disease, *PKD1*, encodes a very large putative membrane protein with an ∼3000 amino acid N-terminus, 11 transmembrane spanning (TM) domains, and an ∼200 amino acid C-terminus ^1^. This enigmatic but essential protein, PC1, undergoes a complex set of post-transcriptional cleavage steps that are incompletely characterized. The best understood occurs at its G-protein coupled receptor cleavage site (GPS) that resides at the boundary of the 3000aa N-terminus and the first TM ^2^. Cleavage occurs in the Golgi, resulting in an N-Terminal Fragment (NTF) that remains tethered to the stalk and remainder of the C-terminal Fragment (CTF). Cleavage at this site is thought essential for proper trafficking to the primary cilium ^3^, and mutations that disrupt cleavage result in severe, distal cystic disease that results in death by around P30 in homozygous mutant mice ^4^.

PC1 has been localized to multiple other subcellular locations, however, including the apical and basolateral membranes and the endoplasmic reticulum ^5-9^, and its cytoplasmic C-terminus has been reported to traffic to the nucleus and to mitochondria ^10-12^. The protein is reported to have multiple functions, including as a flow-sensor ^13^, a Wnt-receptor ^14^, an atypical G-protein coupled receptor ^15^, and has been implicated in cell-cell and cell-matrix signaling ^16, 17^. PC1 cleavage products have been reported to regulate transcription ^10, 11^ and mitochondrial function^12, 18^.

Given PC1’s large size and multiple domains, it is certainly possible that it maps to each of these locations and functions in multiple pathways, but studies have reported conflicting results ^12^. These discrepancies highlight one of the challenges in the field, which is tracking PC1 in an unambiguous function. PC1 expression decreases dramatically in post-natal kidney tissue, and while its expression is higher in other adult tissues, much less work has been done using samples from other organs ^19^. Numerous studies, therefore, have either relied on over-expression of partial or full-length recombinant PC1, often with various epitope tags added to facilitate tracking, but it is not known how well these systems truly recapitulate the endogenous system. Two mouse models have been reported with the HA-epitope either introduced into the endogenous mouse *Pkd1* locus ^20^ or into a Bacterial Artificial Chromosome (BAC) that contains the mouse gene, which was then introduced into the mouse as a transgene ^21^. The advantage of these *in vivo* systems is that they provided direct evidence that introduction of the epitope tags had not seriously altered PC1 function as the mice did not develop cystic disease. However, the HA tag does not allow live cell imaging of PC1 trafficking, and the published studies have described limited detection of the tagged protein in tissues, with all expression limited to the primary cilium.

As noted above, our group had recently discovered that the C-terminus of PC1 (CTT) traffics to the mitochondria using both cell-based imaging and biochemical methods. We also showed that loss of PC1 affected mitochondrial membrane potential and mitochondrial network structure while re-expression of CTT rescued the latter ^12^. A limitation of these studies is that they were performed using recombinant fluorescently tagged *Pkd1* constructs in cell culture systems. While we could also detect CTT in mitochondria in mouse embryonic fibroblasts (MEFs) isolated from transgenic mice carrying the *Pkd1* BAC using biochemical methods, we were unable to unambiguously detect it using imaging methods.

It is important to note that our results differed from prior reports which described a nuclear pattern for a similar C-terminal fragment ^10-12^. Our results also differed from those of another group that had reported a direct link between PC1 and mitochondria but which was restricted to mitochondrial associated membranes (MAMs), contact sites between the ER and mitochondria, rather than internal to mitochondria ^18^. In this schema, the C-terminus of PC1 would presumably be engaging with mitochondrial surface proteins, possibly regulating mitochondrial calcium flux. The two distinct patterns, therefore, suggest different functions for PC1.

Given the many conflicting results for PC1 and the general inability of the PKD community to visualize endogenous PC1, we sought to generate a new mouse line that had an enhanced green fluorescent protein (eGFP) “knocked-into” the C-terminus of PC1 that could be used for both visualizing endogenous PC1 and identifying its endogenous interactome using magnetic bead-linked GFP nanobodies. We also added a triple HA-epitope tag to allow dual purification/tracking of the endogenous protein using antibody-based detection systems. We report here that this modification had minimal effects on endogenous PC1 function but that the endogenous level of PC1 expression is below the threshold of detection in living cells. We were, however, able to use the model to begin to characterize the *in vivo* interactome of PC1 and have identified the mitochondrial protein nicotinamide nucleotide transhydrogenase (Nnt) as a putative binding partner.

## Methods

### Ethics Statement

All mouse studies were performed using protocols approved by NIH Animal Care as appropriate. Mice were kept and cared in pathogen-free animal facilities accredited by the American Association for the Accreditation of Laboratory Animal Care and meet federal (NIH) guidelines for the humane and appropriate care of laboratory animal.

### Pkd1 knock-in model

*Pkd1-GFP* knock-in mice were generated using the CRISPR/Cas9 technology. Briefly one sgRNA (GGTCCACCCCAGCAGCACTT) targeting the last exon of *Pkd1* was designed to cut near the stop codon. The donor DNA construct was made by flanking the eGFP coding region, followed by 3-HA tags, with 5’ (2992 bp) and 3’ (3037bp) homologous arms of the cut site, introducing a silent mutation in the last threonine codon from ACT to ACA (to prevent the donor plasmid from being targeted by Cas9) and then inserting the GFP-3HA between the last codon and the stop codon (TAG). The donor DNA construct (10ng/ul) was co-microinjected with Cas9 mRNA (50ng/ul) and sgRNA (20ng/ug) into the pronuclei of fertilized eggs collected from B6D2F1/J mice (JAX #100006). The injected embryos were cultured overnight in M16 medium overnight, and those embryos which reached 2-cell stage of development were implanted into the oviducts of pseudo-pregnant foster mothers (CD-1 mice from Charles River Laboratory). Offspring born to the foster mothers were genotyped by PCR. Two male mice with confirmed insertion were bred: one was a chimeric with likely no germline insertion, and the other produced pups carrying the knock-in and was therefore our founder mouse. Genomic DNA of the founder mouse was used for long-range PCR of regions flanking the homology arms, GFP-3HA sequence and all exons in the region, which were sequenced and confirmed to be without mutations (supplementary file 1). This animal was bred with C57BL/6 mice to establish the knock-in mouse line. F1 mice were either back-crossed to C57BL/6 mice (up to F4) or interbred. Subsequent genotyping of this line was done with the primers: E46-F: 5’-TGCTTGTCCAGTTTGACCGA with eGFP-R: 5’-GCTGAACTTGTGGCCGTTTA and 3UTR-R: 5’-ATGGCCACCTAGGGGTAGAG (wild type product: 614 bp; knock-in product: 332 bp).

### Mouse studies

*Pkd1*^*ko*^ mice were previously described ^22^. Mice homozygous for the *Pkd1*^*ko*^ allele lack exons 2 to 4 and die *in utero. Pkd1*^*cko*^ mice have loxP inserted into introns 1 and 4, and generates the deleted allele *Pkd1*^*del2-4*^ upon cre-expression ^22^. Tamoxifen-inducible Cre-ER transgenic mice (JAX strain #004682 ^23^) and Ksp-cre transgenic mice (JAX strain #012237 ^24^) expressing cre recombinase in kidney tubular epithelial cells were used to induce *Pkd1* inactivation in kidney. In the Cre-ER line, mice were induced at post-natal day 40 (P40) with 0.4mg/g tamoxifen (Sigma, T5648) diluted 20mg/ml in corn oil and injected intra-peritoneally.

C57BL/6J (JAX strain #000664) and C57BL/6NJ (JAX strain #005304) strains were used as *Nnt* mutant and wild type lines, respectively. *Nnt* mutation was confirmed by PCR as described in ^25^. Briefly, a PCR reaction identifies a 312bp product only in *Nnt* wild type mice using primers: Exon6_L1: CAATTCTGCCAACAACTGGA and Exon6_R4: GGTCACTCTGGGCACTGTTT; and a 547bp product only in mutants using Exon12_L1: GTAGGGCCAACTGTTTCTGC and Exon6_L4: TCCCCTCCCTTCCATTTAGT.

For NAC intervention studies, a cohort of *Pkd1* conditional; Ksp-cre transgenic mice in both *Nnt* mutant or wild type backgrounds was constantly treated with N-acetyl-L-cysteine (NAC; A7250 Sigma) mixed in drinking water at 7 mg/mL ^26^ supplemented with 5 mg/ml of Splenda sweetener, neutralized to pH 7.4 with NaOH and freshly prepared on alternate days and continued during pregnancy and lactation. A corresponding control group of littermate breeding pairs was treated in the same manner with water supplemented with 5 mg/ml of Splenda. Pups were harvested at post-natal day 9 (P9). Power calculations to detect a 20% increase/decrease in kidney/body weight were used to determine number of mice needed in the study. The breeding strategy was consistent with ¼ of the mice being mutant (*Pkd1*^*cko/cko*^*;Ksp-cre+*) and resulted in a total of 32 mutants in NAC, 44 mutants in Splenda, 98 controls in NAC and 89 controls in Splenda. P8 mice were euthanized, kidney and body weights were measured, one kidney was snap frozen in liquid nitrogen and the other was fixed in 10% formalin.

### Primary cell lines

Primary tubular epithelial cells (PTEC) were obtained from kidneys removed from P2 pups. Briefly, kidneys were quickly dissected after mouse euthanization and transferred to gentleMACS C Tubes (Miltenyi Biotec Inc., cat. no. # 130-093-237) in 2ml of digestion buffer (13.5 ml of Dispase (STEMCELL, cat. # 07912) with 1.5 ml of Collagenase/Hyaluronidase 10X (STEMCELL cat. # 0793), 150 µl of GlutaMax, 150 µl of 1M HEPES and 0.75 µl of 10 mg/ml DNAse I) and dissociated in gentleMACS Dissociator (Miltenyi Biotec Inc., cat. no. 130-093-235) using the Multi_E_01.01 program, followed by incubation at 37°C for 30 min. under gentle rotation. Dissociated cells were filtered through pre-separation filters (Miltenyi Biotec Inc. cat. no. # 130-095-823) and centrifuged at 150xg for 5 min. and either directly seeded on 10cm cell culture dishes or further enriched with *Lotus Tetragonolobus Lectin* (LTL; Vector labs biotynilated beads, cat. no. B-1325-2) LTL or *Dolichos Biflorus Agglutinin* (DBA; Vector labs biotinylated beads, cat. no. B-1035-5) beads and CELLection Biotin Binder Kit (Invitrogen, cat. no. 11533D). Cells were grown at 37 °C with 5% CO_2_ in low serum media DMEM/F12 media (Life cat. no. 21041-025) with 2% FBS (GEMINI Bio-Products cat. no. 100-106), 1 x Insulin-Transferrin-Selenium (Thermo Fisher Scientific, cat. no. 41400-045), 5 µM dexamethasone (SIGMA, cat. no. D1756), 10 ng/ml EGF (SIGMA, cat. no. SRP3196), 1 nM 3,3’,5-Triiodo-L-thyronine (SIGMA, cat. no. T6397) and 10 mM HEPES (CORNING, cat. no. 25-060-CI).

Mouse embryonic fibroblasts (MEF) were obtained from E13.5 embryos which were chopped coarsely (10–20 times) using a sterile razor blade and digested in 0.25% Trypsin-EDTA for 10 min. in a cell culture incubator (37 °C; 5% CO_2_) followed by dissociation in gentleMACS C Tubes (Miltenyi Biotec Inc., cat. no. # 130-093-237) in a total of 8 ml DMEM media with 10% FBS using gentleMACS Dissociator Program A twice and seeded onto 15 cm tissue culture dishes and grown in DMEM with 10% FBS at 37 °C with 5% CO_2_.

### Immunoblot and immunoprecipitation

Mouse tissue was dissociated using gentleMACS Dissociator (Miltenyi Biotec Inc., cat. no. 130-093-235) and gentleMACS M Tubes (Miltenyi Biotec Inc., cat. no. # 130-093-236) in lysis buffer (1% Triton X-100; 150 mM NaCl; 50 mM Tris HCl pH8.0, 0.25% sodium deoxycholate, 1 × protease inhibitor with EDTA (Roche Complete, cat. no. 11836153001), 1 x phosphatase inhibitor (Roche PhosSTOP, cat. no. 4906845001), 1 µl/ml Nuclease (Pierce, cat. no. 88700)). Immunoprecipitation of tagged PC1 was done using anti-GFP nanobodies coupled to magnetic agarose beads (GFP-trap, Chromotek cat. no. gtma-10) following the manufacturer’s protocol. For immunoblotting, samples were denatured in 1:1 with 2 x SDS buffer with 1x reducing agent (Invitrogen, NP0004) for 10 min at 95 °C, loaded into 4-12 % bis tris 1.5 mm gels (Invitrogen, NP0336BOX), run in MES buffer (Invitrogen, NP0002) at 200V for 45 min., transferred to nitrocellulose membrane (Invitrogen, IB301002) using iBlot dry blotting transfer system (Invitrogen) program 3 and blocked/incubated with antibodies and imaged using Li-Cor Odyssey Imaging System following the manufacturer’s instructions and the following antibodies: anti-eGFP (Aves, cat. no. GFP-1020), anti-HA (MBL, cat. no. M180-3), anti-Nnt (Thermo Fisher Scientific, cat. no. 13442-2-AP), anti-Polycystin-2 (PC2-CT ^27^, PKD-RCC (https://www.pkd-rrc.org/) and anti-Polycystin-1 (7E12, Santa Cruz cat. no. sc-130554) and secondary labeled antibodies (Licor, cat. no. 925-68072 or 926-32211).

### Mass spectrometry

For each mass spectrometry experiment, 8 heads of P1 newborn mice of each genotype (*Pkd1*^*eGFP/eGFP*^ and *Pkd1*^*wt/wt*^) were used as input for immunoprecipitation (approximately 80mg of protein crude lysate/genotype). Briefly, two heads were combined, dissociated and lysed as described above. Protein concentration was measured using BCA assay (Pierce, cat. no. 23225) and 20mg were used for immunoprecipation (IP) with GFP-trap. The eluates of the four independent IPs for each genotype were then pooled and immediately processed for mass spectrometry. This procedure was repeated three times (using a total of 24 heads). A bottom-up proteomic analysis used a soap to assist in rendering and digestion of sample proteins ^28, 29^, post-digestion reductive demethylation ^30, 31^, neutral pH off line reversed phase separation using fraction concatenation ^32^ conventional High-Low microscale HPLC/mass spectrometry, and analysis using MaxQuant ^33^ and R ^34^ using the ggplot2 package ^35^.

### Histology and immunostaining

Cells were seeded at a density of 100,000 cells/well in chamber slides (Ibidi, cat. no. 80822). After 24h, cells were serum-starved for 48h, fixed for 10 min. in 10% formalin, rinsed with PBS and permeabilized with 0.5% Triton X-100 in PBS for 5 min., blocked with 1% fish gelatin in PBS, stained with antibodies to detect Arl13 (ciliary marker; ProteinTech cat. no. 17711-1-AP; secondary antibody: Invitrogen A27034 labeled with Alexa Fluor 488), γ-tubulin (ciliary basal body marker; SIGMA cat. no. T5192; secondary antibody Invitrogen A27034 labeled with Alexa Fluor 488) or eGFP (Aves cat. no. GFP-1020; secondary antibody: Invitrogen A21449 labeled with Alexa Fluor 647), mounted (Invitrogen ProLong cat. no. P36981) and imaged using Zeiss LSM700 confocal microscope. Freshly dissected embryos or kidneys were frozen in OCT (Tissue-Tek; Sakura Finetek USA) and frozen sections were processed as above; or embedded in paraffin, sectioned, and stained with trichrome staining or used for immunohistochemistry. Images were visualized using Imaris (Bitplane). Cystic index was calculated by delineating kidneys in trichrome stained slides using the magnetic lasso feature in Photoshop, and measuring total kidney area, thresholding cystic area and measuring total kidney area and cystic area using Fiji ^36^.

### Biochemical assays

Abcam assay kits were used to measure NADP+/NADPH (ab65349) and NAD+/NADH (ab65348) following the manufacturer’s instructions. Briefly, frozen kidneys were thawed on ice and homogenized using gentleMACS Dissociator (Miltenyi Biotec Inc., cat. no. 130-093-235) and gentleMACS M Tubes (Miltenyi Biotec Inc., cat. no. # 130-093-236) in Abcam extraction buffer with 1% bezonuclease at 50 mg of tissues/500 ul and filtered through 10 kD spin columns by centrifugation for 15min at 10,000xg. Protein concentration of the lysates was measured using the BCA assay (Pierce, cat. no. 23225) and filtered lysates were analyzed using a plate reader (BMG Labtech CLARIOstar microplate reader) at OD450nm.

### Statistics

The R environment ^34^ was used for statistical analysis. To minimize the effect of outliers, robust statistical estimators using the WRS2 package were used ^37^. In particular, comparisons of two groups were done using the function pb2gen for t-test based on medians. Plots were made using the ggplot2 package ^35^. The pedigree tree network was generated in Cytoscape ^38^.

## Results

### Generation and characterization of Pkd1 knock-in model

Using CRISPR/Cas9 technology, the last exon of *Pkd1* was targeted to insert an enhanced green fluorescent protein (eGFP) followed by three human influenza hemagglutinin (HA) tags in frame with the C-terminus of the protein (Figure 1A-C). Screening of targeted mice revealed one mouse with germline knock-in carrying this expected *Pkd1*^*eGFP*^ allele, confirmed by long-range PCR and RT-PCR (Figure 1D-E). This animal carried a germline *Pkd1*^*eGFP*^ allele and was the founder of the *Pkd1*^*eGFP*^ knock-in line (Figure 1F).

**Figure 1.**
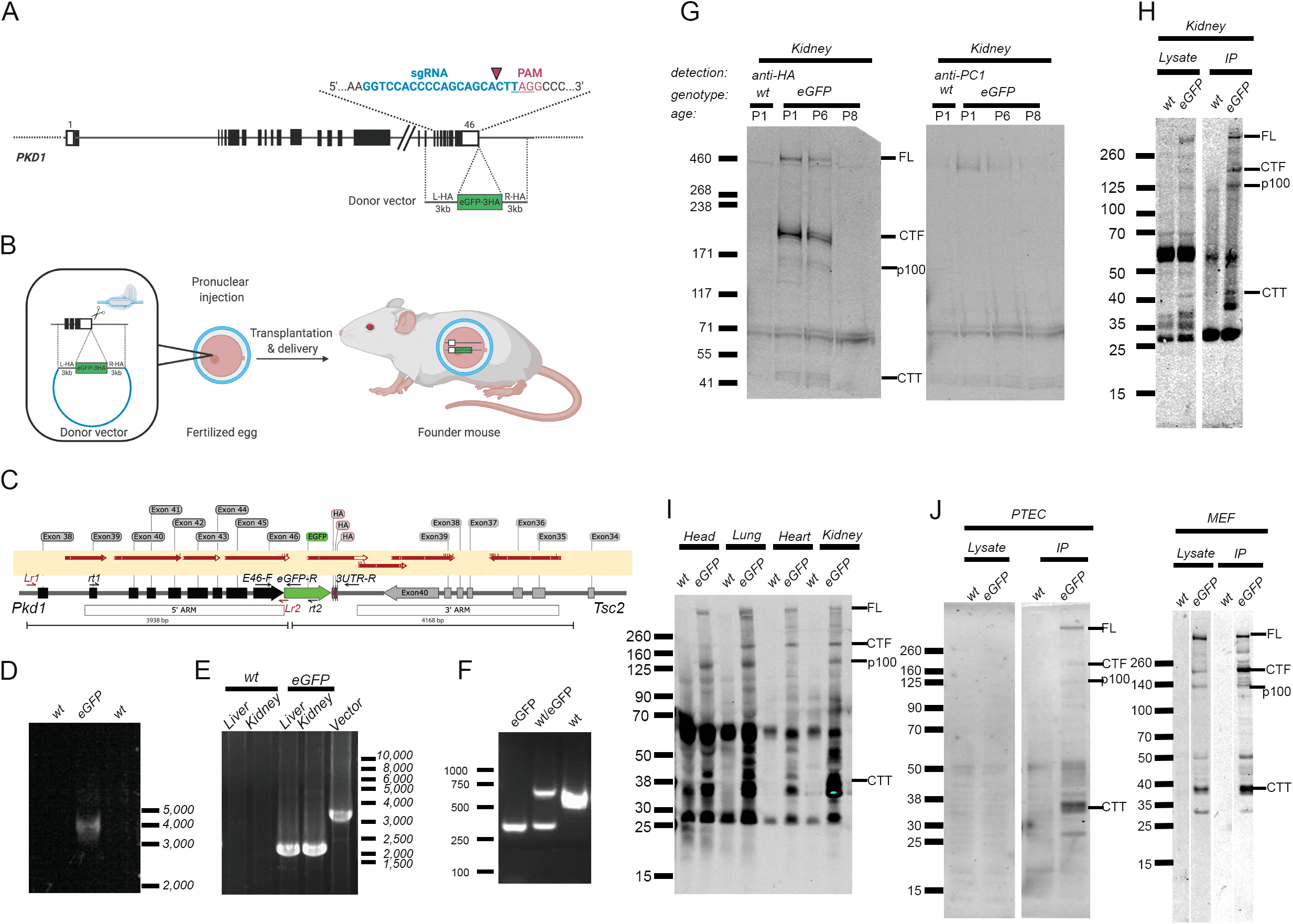
**A)** Schematics of *Pkd1* knock-in allele, showing sgRNA guide sequence and location of eGFP-3xHA tag. **B)** Diagram showing steps to generate the knock-in mouse. **C)** Structure of the knock-in *Pkd1* allele. Red arrows on the light yellow rectangle mark regions confirmed by sequencing. Small arrows in the gene diagram are the locations of primers used in the reactions in panels D-F. **D)** Genomic long-range PCR using primers Lr1 and the eGFP-specific Lr2 showing the predicted band of ∼4kb in *Pkd1*^*eGFP*^ founder mouse. No bands are amplified in the control tissue (wt). **E)** Reverse-transcriptase PCR using primers rt1 and rt2 showing expression of the *Pkd1* transcript with eGFP sequence in knock-in mouse tissue. The vector positive control is larger because it includes introns. **F)** Genomic PCR using primers E46-F, eGFP-R and 3UTR-R to genotype knock-in mice. As expected, eGFP homozygotes only have the ∼300bp E46-F:eGFP-R product, control specimens (wt) have only the 600bp E46-F:3UTR-R product, and heterozygotes have both. **G)** Detection of PC-1 in kidney of wild type and knock-in mice at different post-natal days; left: detected with anti-HA antibody; right: detected with anti-PC1 (7e12) antibody. Full length, uncleaved PC1 is detected with the anti-HA antibody (left) whereas 7e12 only detected NTF under these conditions (right). **H)** Kidney of control and knock-in P1 mice showing tagged PC1 detection with anti-HA antibody; left: cell lysate; right: protein immunoprecipated with anti-eGFP conjugated agarose beads. Note the enrichment of CTT after IP. **I)** Immunoprecipitation of PC1 from different tissues of P1 control and knock-in mice using anti-eGFP conjugated agarose beads and detected with anti-HA antibodies. **J)** Mouse primary tubular epithelial cells (PTEC) and embryonic fibroblast (MEF) cells from control and knock-in mice showing tagged PC1 in the lysate (left) and immunoprecipitated by anti-eGFP and detected with anti-HA (right).

Immunoblot analyses showed that tagged polycystin-1 (PC1-eGFP) is expressed in kidneys of young pups, with highest levels on day 1 and decreasing to barely detectable levels by post-natal day 8 (Figure 1G). In mouse tissue, detection using the tag is more sensitive than using antibodies to the endogenous protein (Figure 1G). Using nanobodies against eGFP, PC1-eGFP can be efficiently enriched in pull-down experiments (Figure 1H-J) and can be detected in multiple mouse tissues, including brain, lung, heart and kidney; and in primary cell lines obtained from knock-in mice.

The founder mouse was bred to wild type and to heterozygous knock-out (*Pkd1*^*ko/wt*^) mice (Figure 2A). *Pkd1*^*eGFP/wt*^ bred to *Pkd1*^*ko/wt*^ produced pups at the expected mendelian rates (Figure 2B). A cohort of 6 *Pkd1*^*eGFP/ko*^ mice were aged to 15 months and showed no kidney pathology, though one animal developed small liver cysts (Figures 2C-E). *Pkd1*^*eGFP/eGFP*^ mice euthanized at multiple ages between birth and 450 days of age had no obvious phenotypes (>10 mice/30-day brackets). These data suggest that a C-terminal tag added to PC1 does not significantly impair its function.

**Figure 2.**
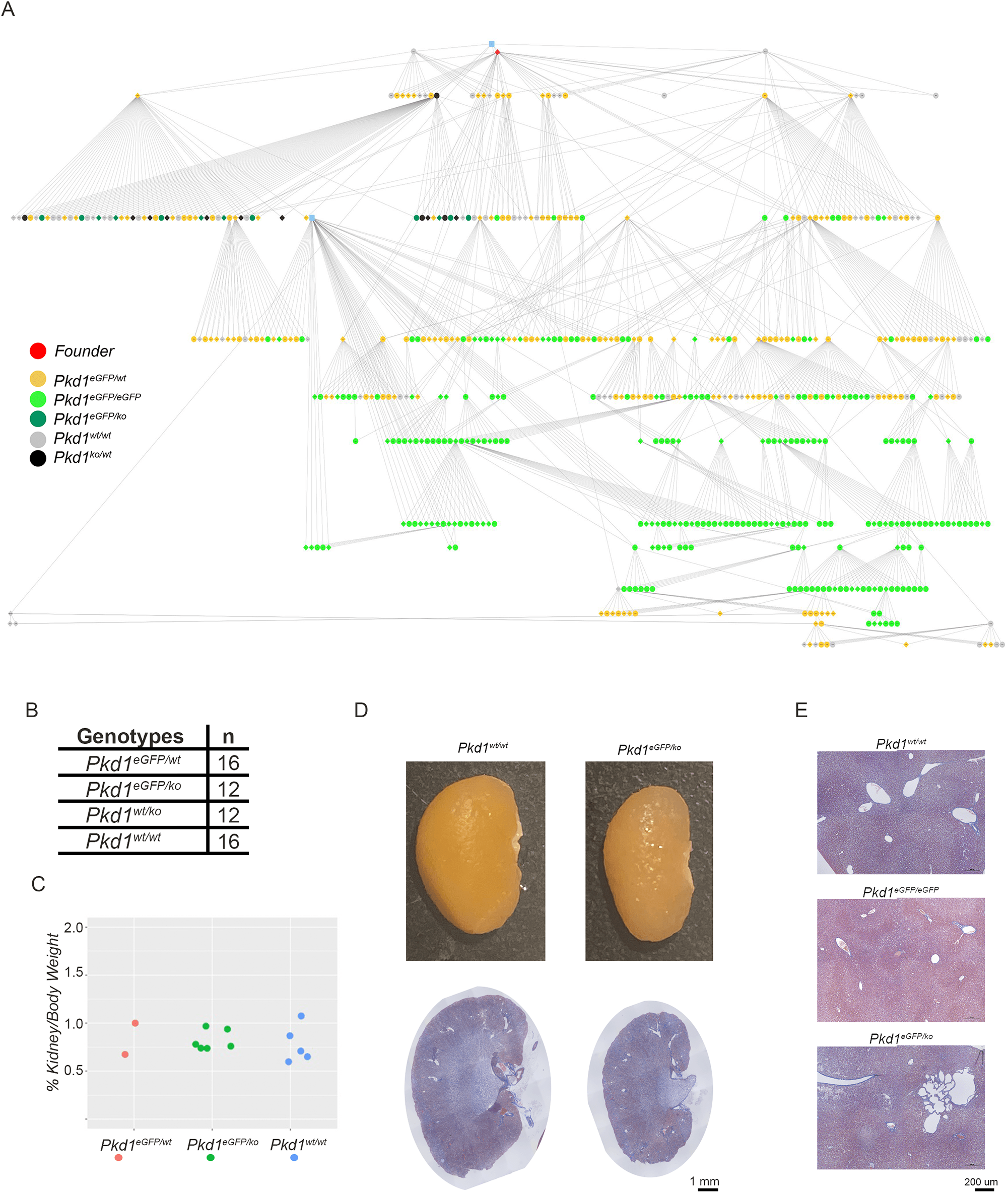
**A)** Pedigree tree of mouse colony derived from the single *Pkd1*^*eGFP*^ knock-in mouse (red dot at the top). Genotypes are color-coded. The blue dots represent wild type mice used for back-crossing. **B)** Table showing that *Pkd1*^*eGFP/ko*^ pups born to *Pkd1*^*eGFP/wt*^ vs. *Pkd1*^*ko/wt*^ breedings are viable. **C)** Kidney/body weight ratios of a subset of animals in panel B allowed to age to 15 months showing normal kidney size. **D)** Representative *Pkd1*^*eGFP/ko*^ kidney has normal morphology at 15 months of age (lower panels: trichrome staining; size bar: 1mm). **E)** While the majority of the *Pkd1*^*eGFP/eGFP*^ and *Pkd1*^*eGFP/ko*^ livers had normal morphology, one 15 months-old *Pkd1*^*eGFP/ko*^ mouse had liver cysts (middle panel; trichrome staining; size bar: 200 µm).

One of the goals of creating a *Pkd1* knock-in mouse was to detect PC1 *in vivo* using microscopy tools. So far, we have been unable to unambiguously identify PC1 in live cells/tissue. We also have been unable to unambiguously, consistently detect PC1 above background in fetal or adult tissues using antibody amplification methods. However, using antibody amplification, we have consistently detected PC1 in the primary cilium of kidney epithelial cells (Figure 3A-B). In mouse embryonic fibroblasts, while a few cilia have detectable PC1 expression, most do not (Figure 3C). We also have not been able to detect PC1 above background in other subcellular structures. However, in some structures (for example endoplasmic reticulum and mitochondria) the signal appeared to be of slightly increased intensity in PC1 samples, suggesting that it is possible PC1 could be detected there (data not shown).

**Figure 3.**
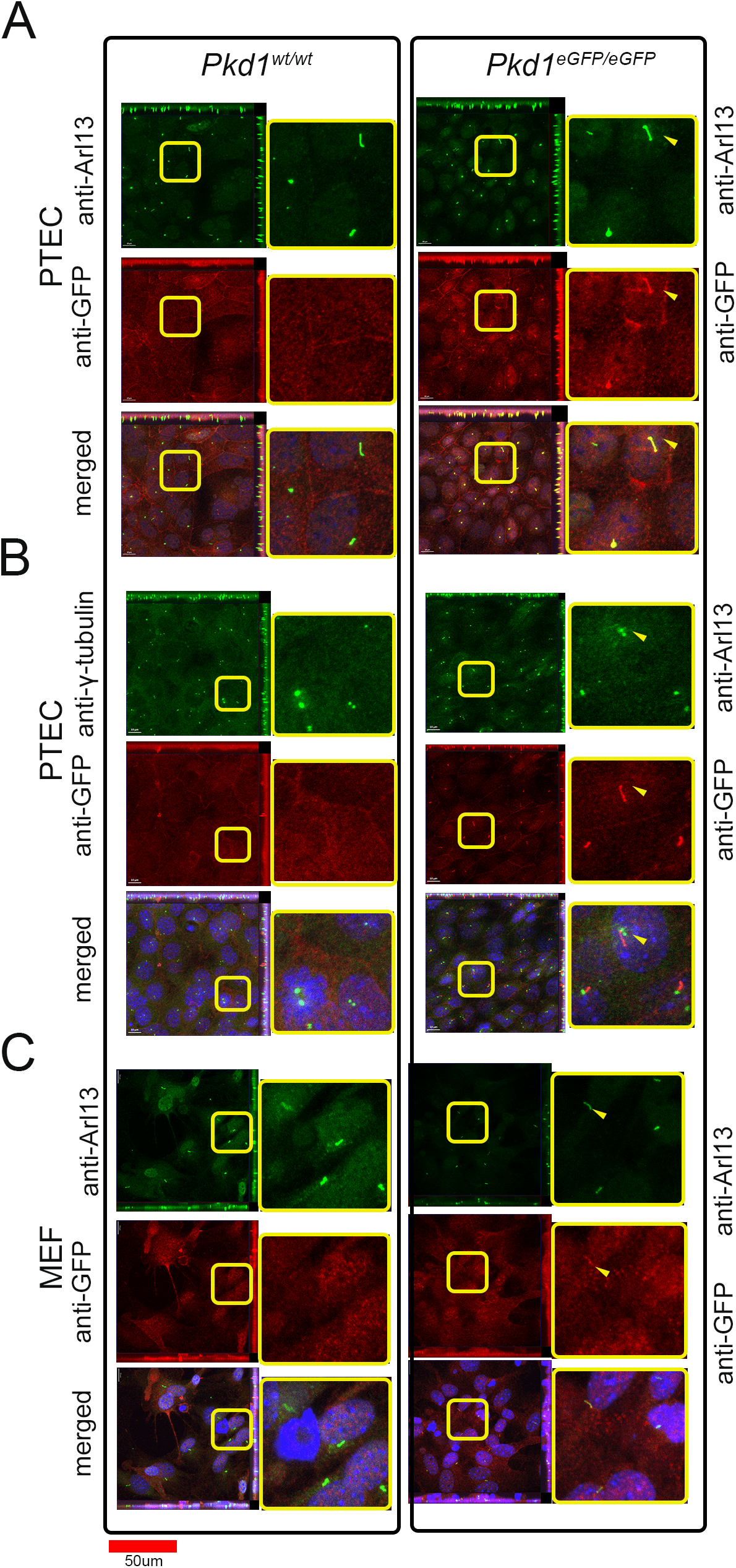
Tagged PC1 can be detected in cilia of primary cells and tissues of knock-in mice. **A)** Detection of PC1 with anti-eGFP antibody in cilia of most primary tubular epithelial cells (PTEC), co-localized with Arl13; **B)** PC1 localizes adjacent to ciliary bodies, marked with anti-γ-tubulin. **C)** PC1 is detected in cilia of some, but not all, mouse embryonic fibroblasts (MEF) cells.

### Using the model to study the in vivo PC1-interactome and to identify a mitochondrial binding partner

Another goal of the *Pkd1* knock-in model was to identify *in vivo* PC1 binding partners and ultimately to establish a PC1-interactome. To determine the feasibility of this goal, we initially optimized conditions to immunoprecipitate PC1 and detect both PC1 and PC2, a known PC1 binding partner ^39^ (Figure 4B). Having established that PC1 and PC2 could be co-immunoprecipitated in mouse tissue, we sought to determine conditions to detect PC1 and binding partners by mass spectrometry. After several experiments using different protocols, tissue and starting amounts (data not shown), it became clear that large amounts of input material are necessary for robust PC1 detection. Using lysate of approximately 80mg of tissue, PC1 and PC2 were invariably detected, and served as a positive control. As proof-of-principle that such strategy could yield meaningful interactors, we performed 3 sets of affinity purification mass spectrometry experiments, each analyzing the immunoprecipitated of combined heads of 8 P1 mice (brain had one of the highest expression levels of PC1 at this age) in both knock-in and control mice (Figure 4). As expected, under these conditions, both PC1 (which was the bait) and PC2 (which forms a complex with PC1) were the proteins with highest enrichment and intensity in knock-in mouse samples (Figure 4C).

**Figure 4.**
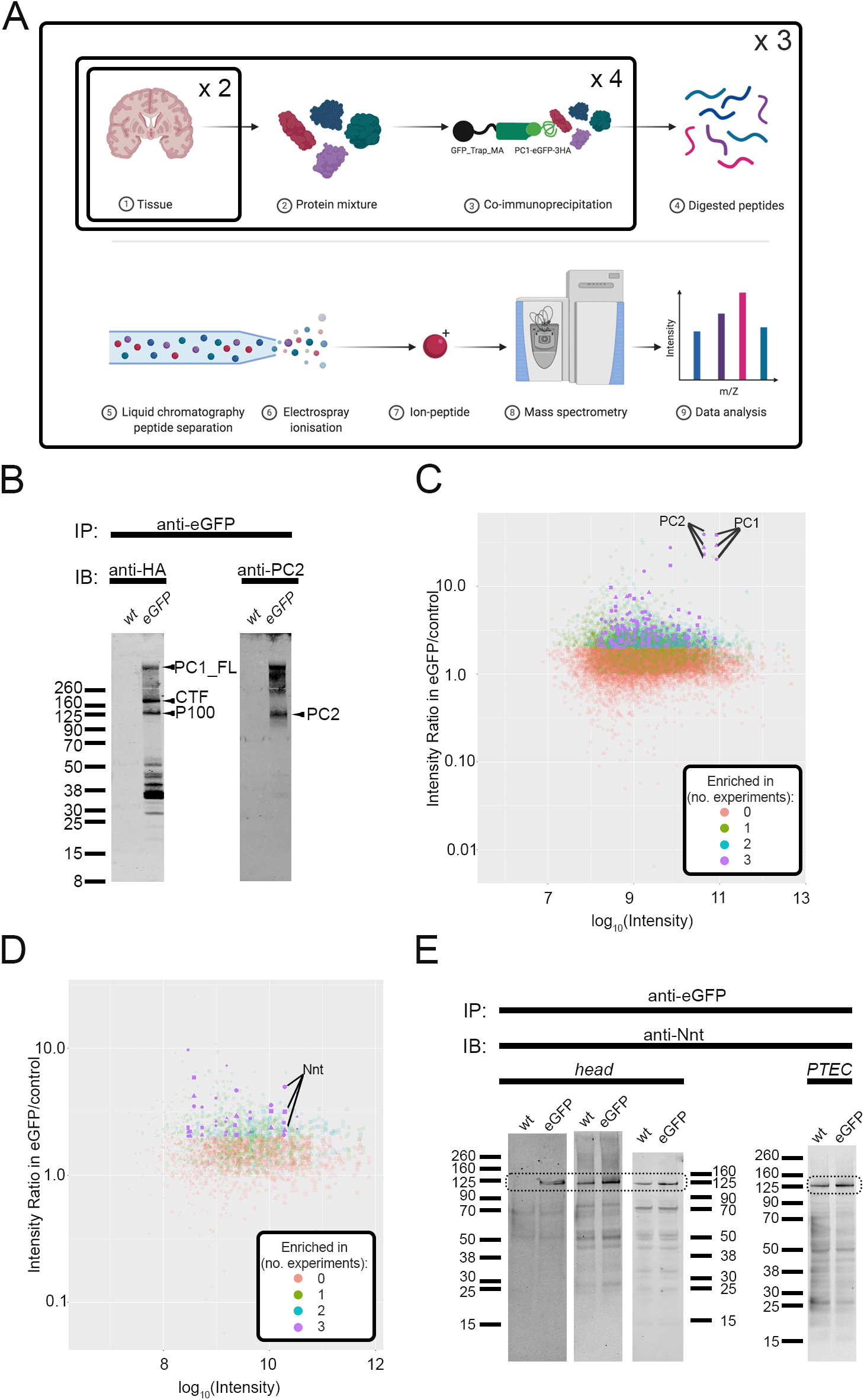
Identification of PC1 *in vivo* binding partners. **A)** Schematics showing steps for proteomics experiments. Briefly, for knock-in and control samples, 4 sets of 2 heads of P1 mice per genotype were independently immunoprecipitated using eGFP beads then pooled and processed for one mass spectrometry experiment. Each mass spectrometry experiment was done three independent times. **B)** PC1 immunoprecipitation in post-natal day 1 (P1) whole mouse body showing on the left panel PC1 full-length (PC1_FL) and two cleavage products (PC1_CT; P100) and on the right panel the same blot, probed with anti-PC2 and detected in a different wavelength, showing PC2 co-immunoprecipitation. **C and D)** Plot showing the intensity (x-axis) and intensity ratio between knock-in and control samples (y-axis) prepared from P1 mouse heads. Each dot corresponds to one detected protein; shape (circle, square, triangle) represents experimental batch; the colors summarize the number of times the enrichment ratio was above 2 for the protein. Proteins that were never enriched in the knock-in IP are orange; proteins enriched in all three experiments are colored purple. In C) all detected proteins are shown and in D) only proteins localized to the mitochondrion (gene ontology category GO:0005739). Panel C shows that PC1 and its primary binding partner PC2 were reproducibly the top hits in each IP study. **E)** Immunoblot showing protein immunoprecipitated using anti-eGFP nanobody in control (wt) and *Pkd1*^*eGFP/eGFP*^ heads and primary tubular (kidney) epithelial cells (PTEC) and detected with anti-Nnt antibodies. The doted box indicates the region with Nnt bands.

When we focused on detected proteins that were previously reported to localize to mitochondria, several were enriched in the knock-in mouse, suggesting they might be part of the mitochondrial PC1 interactome. Among these, nicotinamide nucleotide transhydrogenase (Nnt) was enriched in all replicates (Figure 4D). We confirmed by immunoblot that Nnt was enriched in the PC1-immunoprecipitates of both brain and primary tubular epithelial cells of the knock-in mice, compared to controls (Figure 4E).

### Evidence for genetic interaction between PC1 and Nnt

Nnt is an inner mitochondrial membrane protein that uses H^+^ reentry into the mitochondrial matrix and NADH to reduce NADP^+^ into NADPH, thereby increasing NADPH, the “universal cellular reduction currency” in cells ^40^, which in turn contributes to H_2_O_2_ detoxification ^41^ (Figure 5A). A spontaneous *Nnt* mutation resulting in exon 1 missense and multiexon deletion is present in the C57BL/6J mouse strain and severely compromises Nnt function ^25, 42^. To investigate if a PC1-Nnt interaction could have functional consequences, we bred two *Pkd1* conditional mouse lines (Ksp-cre: an early-onset model of severe cystic predominantly in distal nephron segments; and cre-ER, a tamoxifen-inducible model, induced at P40) ^22-24^ to either C57BL/6J (*Nnt* mutant) or C57BL/6NJ (*Nnt* wild type) and monitored if *Nnt* status correlates with kidney cystic disease severity. In the Ksp-cre model, at age 8 (P8) *Pkd1* mutants show highly cystic kidneys (as measured by 2 kidneys/body weight ratio; KBW), and *Nnt* wild type and mutant strains showed no significant difference, albeit disease severity was more variable in the *Nnt* wild type background (Figure 5B; n=10 *Nnt* mutants and 34 controls). Using the cre-ER model, we induced mice at P40 and harvested them between P180 and P200 and again observed no effect of *Nnt* status on KBW (Figure 5C; n=6-7 per group).

**Figure 5.**
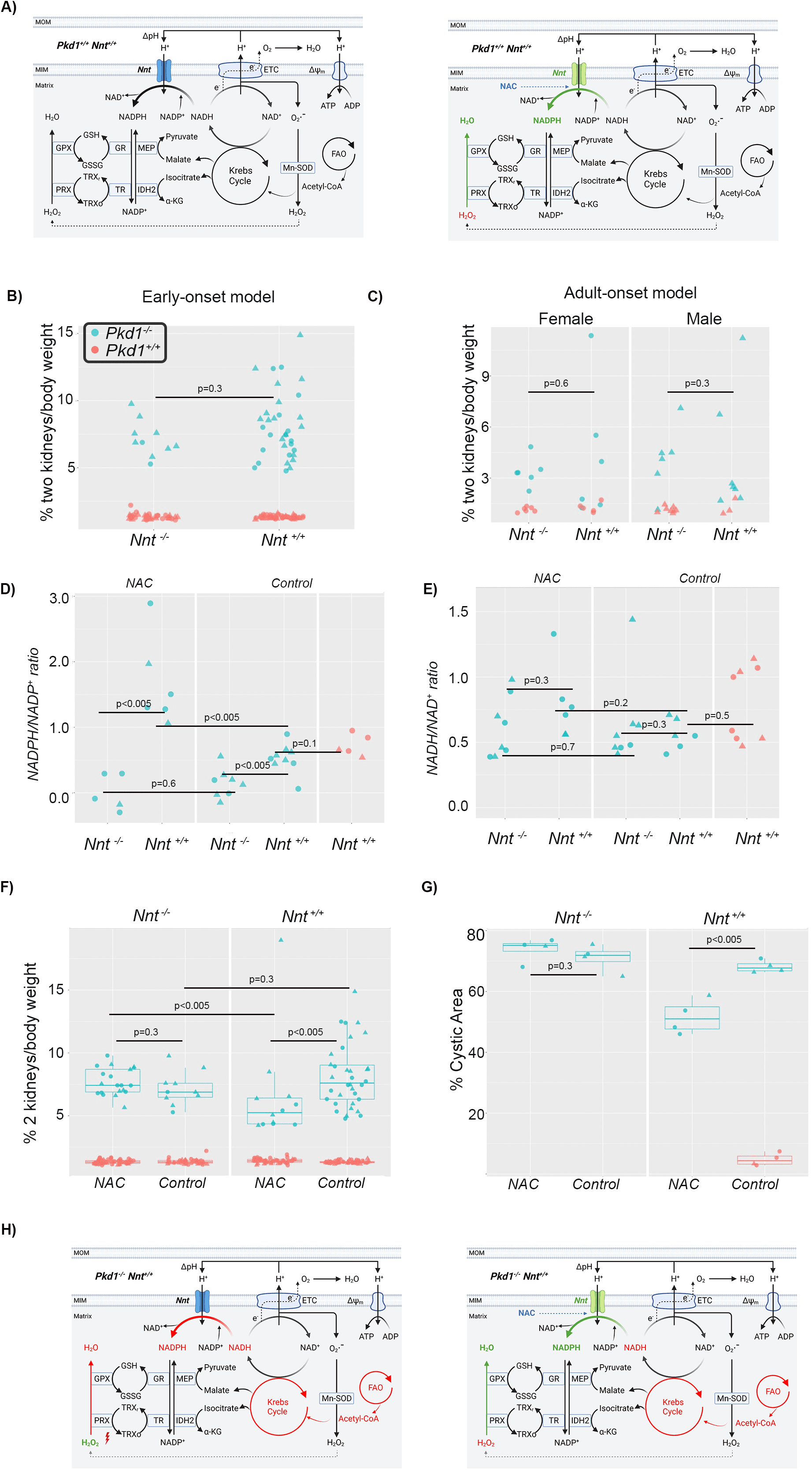
*Nnt* status and PKD progression. **A)** Diagram showing Nnt in the inner mitochondrial membrane. The panel on the left shows baseline activity, and the panel on the right shows presumed changes in the presence of N-acetyl-cysteine (NAC); (green: increased; red: decreased) **B and C)** *Nnt* status does not correlate with kidney/body weight. Kidney/body weight in *Pkd1* mutants (blue) and controls (red) by *Nnt* status in P8 mice induced with nephron-specific Ksp-cre (B; n= 10 *Pkd1/Nnt* mutants and 34 *Pkd1* mutant/Nnt controls) or induced with tamoxifen at P40 and harvested between P180-P200 (n=6-7 of each sex/group). Sex of each mouse is identified by its shape (triangles: males; circles: females). The mice in B) were in a control cohort drinking water with sweetener. **D)** NADPH/NADP+ ratio is decreased in Nnt mutants. The graph shows NADPH/NADP+ ratio in various *Pkd1*/*Nnt* genotypes and treatment conditions. (NAC: N-acetyl-cysteine treated water; control: control water treated with sweetener (NAC treatment: n=5 *Pkd1* mutant; *Nnt-/-;* n=6 *Pkd1* mutant; *Nnt+/+*; control treatment: n = 8 *Pkd1* mutant; *Nnt-/-*; n=9 *Pkd1* mutant; *Nnt+/+*; n= 5 *Pkd1+/+*). Sex of each mouse is identified by its shape (triangles: males; circles: females). **E)** NADH/NAD+ ratio is not significantly changed. The graph shows NADH/NAD+ ratio in various *Pkd1, Nnt* and treatment conditions. (NAC: N-acetyl-cysteine treated water; control: control water treated with sweetener (NAC treatment: n=8 *Pkd1* mutant; *Nnt-/*-; n=6 *Pkd1* mutant; *Nnt+/+*; control treatment: n = 8 *Pkd1* mutant; *Nnt-/-*; n=6 *Pkd1* mutant; *Nnt+/+*; n= 8 *Pkd1+/+*). Sex of each mouse is identified by its shape (triangles: males; circles: females). **F and G)** NAC treatment decreases cystic burden in *Pkd1* mutants in *Nnt+/+* background. In F), the graph shows % (2 kidneys)/body weights in *Pkd1* mutants in either *Nnt* mutant (left panel) or wild type (right panel) backgrounds treated with NAC or control (Splenda without NAC). The *Pkd1-/-* mice in the control-treated group are the same animals shown in panel B (n= 20 *Pkd1-/-;Nnt-/-;*NAC; n= 10 *Pkd1-/-;Nnt-/-;*control; n= 12 *Pkd1-/-;Nnt+/+;*NAC; n= 34 *Pkd1-/-;Nnt+/+;*control). In G), the graph shows the % of cystic area in trichrome-stained kidney sections of a randomly selected subset of mice shown in **F**). **H)** Diagram showing presumed metabolic changes in *Pkd1-/-;Nnt+/+*. Impaired fatty acid oxidation (FAO) and reduced Krebs cycle activity might result in decreased NADPH levels and decreased detoxification of H_2_O_2_ (left panel). With NAC treatment, NADPH levels are increased, resulting in reduced oxidative stress and cystic disease (right panel; green: increased; red: decreased).

Previous reports have suggested mitochondrial dysfunction in PKD ^18, 43, 44^. We therefore wondered if reactive oxygen species (ROS) could play a role and, if so, if reducing ROS damage could: a) slow kidney disease progression; b) be mediated by Nnt. To test this, we treated *Pkd1* mutants with the antioxidant N-acetyl-L-cysteine (NAC), a synthetic precursor of glutathione (GSH) ^45^. GSH is a scavenging antioxidant that forms one of the main lines of defense against reactive species in most cells. Furthermore, NADPH supply is reported to be limiting in GSH synthesis ^46, 47^ in certain conditions. Our hypotheses were therefore that NADPH/NADP+ ratios would be lower in *Nnt* mutants (due to reduced NADPH synthesis) and in *Pkd1* mutants (due to increased ROS damage), and that treatment with NAC would increase the ratio (by reducing ROS damage) in *Pkd1* mutants in Nnt wild type background but would have less effect in an *Nnt* mutant background (due to the limiting NADPH supply). These predictions were tested by measuring NADPH/NADP+ ratios in kidneys of P8 Ksp-cre *Pkd1* mice in *Nnt* mutant and wild type backgrounds. Our results showed that *Nnt* mutants indeed have lower NADPH/NADP ratio in either the NAC-treated (p<0.005; n=5 *Nnt-/-* vs. n=6 *Nnt+/+*) or control-treated groups (p<0.005, n=8 *Nnt-/-* and n=9 *Nnt+/+;* Figure 5D). Furthermore, the effect of NAC treatment was only significant in *Nnt+/+* (p<0.005, n=6 *Nnt+/+* treated with NAC and n=9 *Nnt+/+* in control-treatment), suggesting Nnt activity is rate-limiting for NADPH redox capacity. We did not, however, detect robust differences between *Pkd1* mutants and controls (p=0.110; n= 9 *Pkd1* mutants in control treatment; n= 5 *Pkd1+/+* in control treatment).

Metabolic changes have also been reported in PKD and we wondered if they could be responsible for the NADPH/NADP+ ratio differences. We therefore measured the NADH/NAD+ ratio, which is thought to be less correlated with ROS status and Nnt activity. The ratios were not significantly different across comparisons (Figure 5E), suggesting that ROS pathways are responsible for the observed differences in NADPH/NADP+ ratios.

These results prompted us to test whether NAC treatment could have different effects on phenotype depending on *Nnt* status. Using statistical tests based on robust estimators to compare NAC effect in *Pkd1* mutants in *Nnt* wild type and mutant backgrounds ^37^, we found that NAC treatment had no effect on *Pkd1* mutants in the *Nnt* mutant background, consistent with the NADPH/NADP+ ratio results. In contrast, *Pkd1* mutants in the *Nnt* wild type background benefited from NAC treatment, showing significant differences between medians in kidney/body weight or cystic index (Figure 5F-G).

## Discussion

PC1 has been slow to give up its secrets since the initial description of its primary sequence in 1995. Both its function and even its subcellular localization remain incompletely defined with conflicting reports adding to the uncertainty. We reasoned that an unambiguous localization of PC1 and identification of its interactome would provide a sound basis for assessing its function.

Fluorescently tagged proteins have proven extraordinarily useful for determining patterns of cellular expression in living organisms and cells, and frozen and fixed specimens ^48^. They can be used to track the dynamic process of protein movement within a cell, and using tissue clarification methods, can be used to provide a comprehensive picture of the pattern of expression in an organ ^49^. The protein eGFP is a particularly attractive target given its brightness and the availability and development of nanobodies that have very high specificity and affinity that can be used for its purification ^50, 51^.

We therefore used CRISPR/Cas9 ^52^ to add eGFP and three HA epitope tags in-frame to the very end of the C-terminus of PC1. We confirmed that the integration went as predicted by genomic PCR and DNA sequencing and that both the HA tags and eGFP were expressed by RT-PCR. Mice either homozygous for the modified *Pkd1* allele or had it “in trans” with a *Pkd1* null allele were born at expected mendelian frequencies and aged normally without developing renal cystic disease. These findings definitively show that the addition of this relatively long “foreign” protein sequence had minimal effect on PC1 processing or function. It is worth noting that this “lack” of an effect is unlikely to be explained by rapid cleavage and release of eGFP-3HA from the rest of PC1, as is sometimes noted in recombinant systems, as the sequence is present in both full length PC1 and its various normal cleavage products (CTF, P100, CTT, etc). We did observe one aged *Pkd1*^*eGFP/ko*^ mouse with mild cystic liver disease, suggesting that under certain conditions the added sequence might slightly impair PC1 function, but we conclude that in general the new allele functions properly.

One major disappointment was our inability to track PC1 in tissues or cells of mice homozygous for the modified allele without antibody amplification methods despite multiple efforts to do so. Even using amplification methods and well-characterized antibodies that detect HA and GFP with high sensitivity, we had difficulty detecting PC1-eGFP-3HA reliably above background in tissue specimens. These studies strongly suggest that one of two non-mutually exclusive possibilities: 1) PC1 expression is very low and below the threshold of detection above background; and 2) PC1 expression is dispersed broadly so unable to detect about background because of lack of distinct clustering. This property will mean that unambiguous tracking of endogenous PC1 will remain challenging for the foreseeable future. Our studies do suggest, however, that PC1 can easily tolerate the addition of a sizeable polypeptide to its C-terminus without disrupting its function. It is likely, therefore, that alternative labeling methods using enzymes like the HaloTag or SNAP-tag will also be well-tolerated, though it is unknown whether the signal-to-noise ratio will truly be significantly better for these systems.

Knowing what proteins are associated with one’s target can provide useful clues into its functions ^53^. We designed the new *Pkd1* allele with the explicit goal of being able to characterize the endogenous interactome of PC1. We confirmed in pilot studies that the nanobodies for GFP were extremely efficient for purifying PC1, and we then systematically worked through conditions to determine a starting amount of tissue necessary to ensure that PC1 and its known interactor, Polycystin-2, were consistently among the top hits. We reasoned that as a necessary minimal standard the “bait” being used to isolate other proteins should be one of the most abundant proteins identified by mass spectrometry, and we were reassured to see that PC2 was also always identified as a “positive control”.

To achieve this minimal standard, however, required a substantial scale-up of starting material far beyond what is typical for this type of study. This observation further illustrates just how low endogenous PC1 expression is in tissue and why its characterization has been so challenging. For the present study, we chose to use P1 mouse head as the starting material because it is both one of the largest structures in young mice and has one of the highest levels of PC1 expression. This strategy allowed us to minimize the number of mice required for each experiment, but even this approach required samples from eight young mice as starting material for a single IP. We acknowledge that kidney tissue would have been a preferable starting material since there is a clear functional relationship between loss of PC1 expression and a disease phenotype but this would have required many more mice for each experiment. We reasoned that protein targets of interest could be later validated in kidney samples, an assumption which we subsequently confirmed.

Our approach succeeded in reproducibly identifying a moderate number of likely targets, including mitochondrial proteins, which had relatively low “scores” in the Contaminant Repository for Affinity Purification (“CRAPome”) (https://reprint-apms.org) ^54^, the online database that provides a quantitative and semi-quantitative assessment of a putative protein interactor to be “non-specific.” Full characterization of the interactome is beyond the scope of the present work, but we did focus on one target, Nnt, as an exemplar. While cell lysis and homogenization protocols may bring together proteins that would in physiological state never see each other, Nnt is a mitochondrial protein that is localized to the inner mitochondrial membrane with N-and C-termini within the matrix, a cellular localization where we had seen PC1 previously ^55^. If binding to Nnt was validated and shown to be functionally relevant, it would support our prior studies that localized the C-terminus of PC1 to within the mitochondria. Nnt also plays a key role in maintaining a reduced glutathione pool and regulating reactive oxygen species (ROS) levels within the mitochondria ^56^. The latter properties made it particularly appealing given the many reports implicating elevated ROS in the pathobiology of disease ^18, 43, 44^ and our prior finding that renal epithelial cell lines lacking *Pkd1* have increased mitochondria membrane potential (ΔΨm) ^12^. If PC1 positively regulates the function of Nnt, loss of PC1 could reduce Nnt activity, resulting in increased R0S production and increased ΔΨm. We therefore verified that Nnt was a likely interactor using a second detection system. We also confirmed that the interaction could be detected in primary kidney tubular epithelial cells, further validating our approach of using P1 heads as a useful starting material.

While Nnt’s role as the principal regulator of NADPH levels in the mitochondria is well established, the functional consequences of loss of Nnt are less clear. Numerous studies suggest a role for Nnt in regulating systemic metabolism through its role in regulating insulin secretion in beta cells of the pancreas ^57^. Mice lacking Nnt are reported to develop hyperglycemia, insulin resistance, and obesity when placed on a high fat diet ^58^. Nnt is also thought to play an important role in regulating vasodilation in the vascular system. Mice in the C57BL6J background develop a more severe hypertensive phenotype when treated with Angiotensin II ^59^. They also are more sensitive to loss of Mn^2+^-dependent superoxide dismutase, developing a cardiomyopathy that is partially rescued by re-expression of one wild-type Nnt allele ^60^. In contrast, Nickel found that pathological metabolic demand in an overloaded heart results in reversed direction of Nnt activity, resulting in increased ROS production and worse heart failure ^61^. This study suggests that Nnt can run in both directions, possibly either enhancing or lessening ROS production depending on cellular metabolic demands in a tissue-specific manner. Therefore, it was impossible to predict *a priori* whether loss of *Nnt* in the setting of an absence of *Pkd1* in the kidney would worsen or lessen cystic disease. To our surprise, there was no significant difference in the severity of cystic disease in either the early or late onset *Pkd1* mutant models in either C57BL6J or C57BL6N strains. Despite these negative results, treatment with N-acetyl-cysteine, an agent widely used to treat ROS ^45^, only ameliorated renal cystic disease in the setting of functional Nnt. While interesting, the data do not establish a direct functional link between PC1 and Nnt in the kidney, and future studies will be required to resolve this issue.

In this report, we have characterized only one member of the PC1 interactome of mouse head as proof of principle that this new mouse model can be used for this purpose, but many other proteins were similarly reproducibly pulled down by PC1. Preliminary analysis of this proteome identified other mitochondrial and ciliary proteins within the mix. We suggest that further characterization and validation of this proteome will provide additional novel insights into PC1 function.

In sum, we have produced a new model of PC1 with a fluorescent tag knocked in-frame into the C-terminus of PC1 and used it to show that PC1 expression is too low to detect reliably above background using multiple microscopic methods. Efforts to track endogenous PC1 expression will require more sensitive methods that can detect or tags that can provide much more intense signal. This new mouse model can be used, however, to study the PC1 interactome if a sufficiently large amount of starting material is used. We suggest that as a minimal standard all future interactome studies based on IP methods should detect PC1 as one of the top targets if it is being used as the bait. Finally, we have identified Nnt as a probably bona fide binding target of PC1, further supporting a role for PC1 in the regulation of mitochondrial function.

## Acknowledgments

This research was supported by the NIH, National Institute of Diabetes and Digestive and Kidney Diseases (NIDDK) Intramural Research Program, grant 1ZIADK075042.

We thank Jeff Reece, MS, Director of the Advanced Light Microscopy & Image Analysis Core, NIDDK and Dr. Jiji Chen at the Advanced Light Microscopy and Image Analysis Core, National Institute of Biomedical Imaging and Bioengineering (NIBIB) for helpful suggestions for imaging studies; Dr. Chengyu Liu, Director of the Transgenic Core Facility at the National Heart, Lung, and Blood Institute (NHLBI) for help developing the CRISPR/Cas9 knock-in model; Dr. Zu-Xi Yu at the Pathology Facility, NHLBI, for help histological specimens; and Dr. Markus Rinschen, Aarhus University, for useful advice on mass spectrometry studies. Cartoons in figures 4 and 5 were generated using BioRender.com; figure 4A was adapted from “Protein Peptide Mass Spectrometry”, retrieved from https://app.biorender.com/biorender-templates.

